# Signatures Of Tspan8 Variants Associated With Human Metabolic Regulation And Diseases

**DOI:** 10.1101/2020.11.17.386839

**Authors:** Tisham De, Angela Goncalves, Doug Speed, Phillipe Froguel, NFBC consortium, Daniel Gaffney, Michael R. Johnson, Maarjo-Riitta Jarvelin, Lachlan JM Coin

## Abstract

Here, with the example of common copy number variation (CNV) in the TSPAN8 gene, we present an important piece of work in the field of CNV detection, CNV association with complex human traits such as ^1^H NMR metabolomic phenotypes and an example of functional characterization of CNVs among human induced pluripotent stem cells (HipSci). We report TSPAN8 exon 11 as a new locus associated with metabolomic regulation and show that its biology is associated with several metabolic diseases such as type 2 diabetes (T2D), obesity and cancer. Our results further demonstrate the power of multivariate association models over univariate methods and define new metabolomic signatures for several new genomic loci, which can act as a catalyst for new diagnostics and therapeutic approaches.

## Introduction

In human genetics, the concept of ‘common genetic variation in common diseases’ has been the central tenet of research for more than two decades. Some landmark studies in this field included the thousand genomes project ^1^ (preceded by Hapmap project ^2^) & Genome aggregation database^3^ (gnomAD)-aimed at creating a deep catalogue of human germline mutations, the Wellcome Trust Case Control Consortium (WTCCC) study- aimed at understanding the role of common genetic variation in humans (SNPs^4^ & CNVs^5^) in common diseases, the HipSci project^6^-aimed at elucidating the role of common genetic variants in human induced pluripotent stem cells, The Cancer Genome Atlas (TCGA) project^7^-aimed at comprehensively cataloguing common and recurrent somatic mutations in cancer, The Genotype-Tissue Expression (GTEx) project^8^ - to determine common gene expression patterns across different tissues of the human body and more recently The Human Cell Atlas (HCA) project^9^-aimed at finding common gene expression patterns across millions of single cells from the human body. In addition, well established longitudinal studies such as the Northern Finland Birth Cohorts^10,11^ (NFBC) and UK Biobank^12^ (UKBB) are powerful resources for uncovering the effect of common genetic variants on quantitative traits and lifestyle phenotypes such as socio-economic status, medication, diet etc.

Building on the theme of common genetic variants and their role in common diseases and by integrating insights from almost all of current important landmark human genetic resources, our study here exemplifies that common human genetic variation, in particular common CNVs in the TSPAN8 gene, can play an important and common role in the pathogenesis of diabetes, obesity and cancer. Further, these manifestations are most likely caused through metabolic dysregulation. Through in-depth gene expression analysis including from the human induced pluripotent stem cells project (HipSci) and phewas results for TSPAN8, METTL7B (a trans CNV-QTL for TSPAN8) and NKX2-2 (a common transcription factor), we demonstrate plethora of new evidence which strongly suggest that TSPAN8, METTL7B and NKX2-2 are expressed in tandem in different tissues of the body in humans and in other species and are likely to be linked through molecular functions.

## Results

### TSPAN8 CNVs

The Wellcome trust landmark study of eight common diseases first reported a common CNV (CNVR5583.1, TSPAN8 exon 8 deletion) associated with type 2 diabetes^5^. CNVR5583.1 was PCR validated and was found to have an allele frequency of 36% and 40% for cases and controls respectively. One of the best tagging SNPs for CNVR5583.1 was reported to be rs1705261 with r^2^=0.998-highest linkage disequilibrium (LD) amongst all SNPs. CNVR5583.1, in spite of being a common exonic variant for controls (MAF=40%; highest frequency amongst all WTCCC disease and control cohorts), it has since has not been reported or rediscovered by any of the recent large scale CNV discovery projects. These include the thousand genomes project (n=2,504), the gnomAD project(n=141,456) and more recently the CNV analysis from UKBB^13^ (n=472,228).

Keeping these observations in mind, here, we have reported the rediscovery of CNVR5583.1 in the 1KG next generation sequence (NGS) data for multiple human populations including Finnish (FIN) and British (GBR) populations. Using cnvHitSeq (see methods) we report the CNV deletion frequency for CNVR5583.1 in FIN and GBR as 37.7% and 30% respectively (**Figure 1b**). Next, guided NGS derived breakpoint information (for common exonic CNVs in 1KG) and SNP tagging CNV information (LD) from the WTCCC 16K CNV study, we identified known and novel CNVs in TSPAN8 exon 10 and TSPAN8 exon 11 in two Northern Finland population cohorts- NFBC 1986 (n=4,060) and NFBC 1966 (n=5,240) (**Figure 1a)**. In NFBC 1986, genotyped on Illumina Cardio-Metabochip platform^14^, we rediscovered CNVR5583.1 with an allele frequency of ~2% (**Supplementary Table ST 2d**) tagged by rs1705261 with r^2^= 0.974 (**Supplementary Table ST 2a**). In addition we discovered a new common CNV (MAF~5% in NFBC 1986 and 1KG FIN) overlapping with exon 11 in TSPAN8 which was found to be in weak LD with CNVR5583.1. The LD results were SNP-CNV r^2^=0.623 (**Supplementary Table 2a**) and CNV-CNV r^2^=0.68 (**Extended data figure 9b, 9c**). We highlight that in the public release of CNV data from gnomAD consortium, five common CNVs with MAF>5% were reported in TSPAN8 but none of these were exonic or overlapped with CNVR5583.1 or TSPAN8 exon 11 (**Supplementary Table 2e**). Though we find that there are marked visual differences in sequencing depth coverage across TSPAN8 exon 11 and exon 8 (CNVR5583.1), indicating the presence of structural variation in these regions (**Supplementary Table 2n**). The 1KG CNV release reported no common CNVs within the TSPAN8 gene (**Supplementary Table 2f**). In the TGGA project data, consisting of ~87,000 samples across 287 different cancer types, we observed that TSPAN8 common germline deletions, including CNVR5583.1, are almost completely depleted (VAF<0.01%). In contrast, in most cancer types where TSPAN8 was found to be altered ~2% of the 87,823 patients (91,339 samples from 287 studies), most patient genomes had amplifications with VAF>5% (**Extended data figure 8a**). CNV analysis of the HipSci patient germline genomes and the donor-derived cell lines data indicated a similar pattern. We found TSPAN8 CNV deletions with MAF=5% in germline genomes (**Extended data figure 7c**) and this was reduced to VAF<0.01% in the patient derived iPS cell lines.

**Figure 1.**
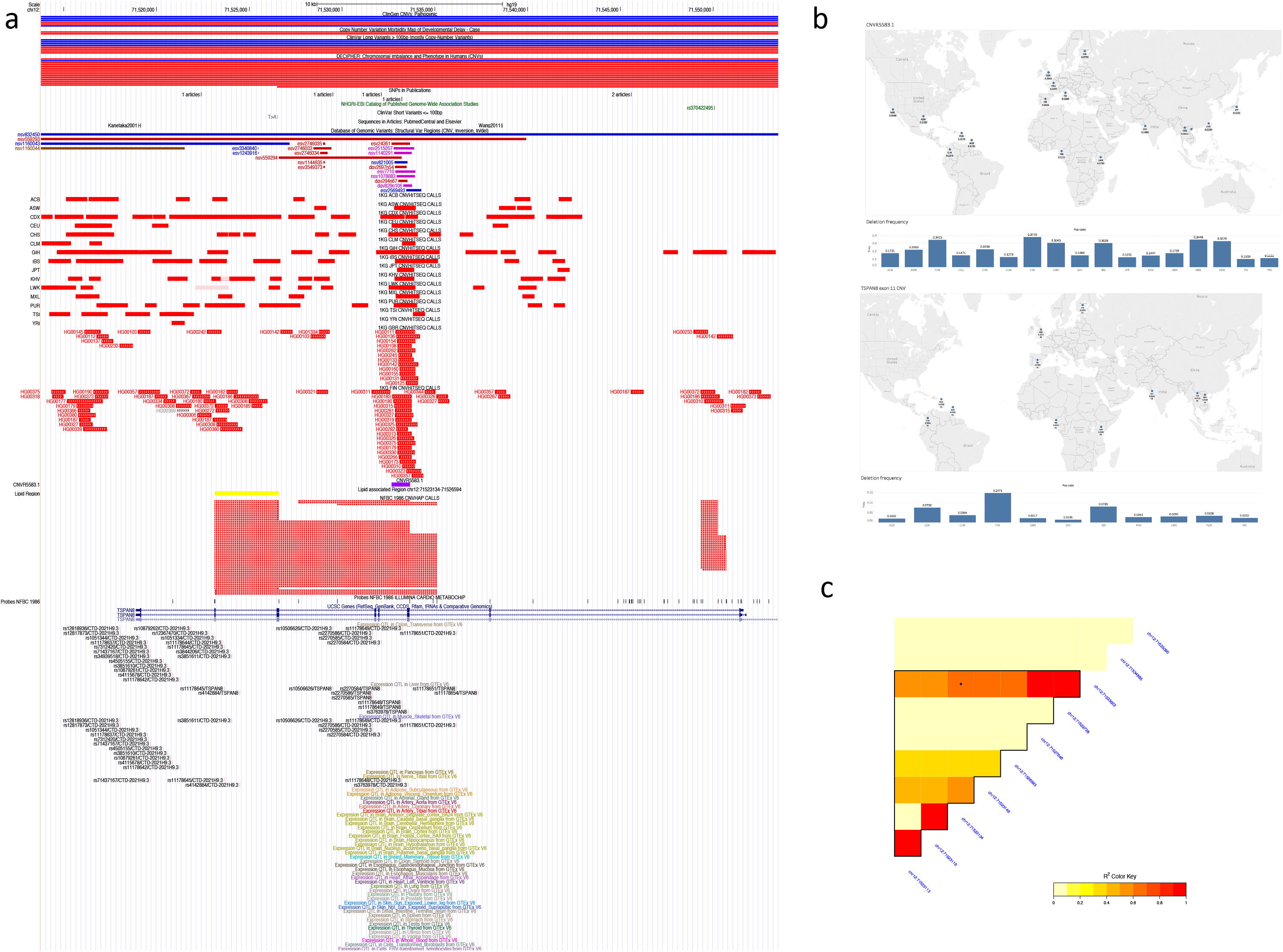
**a** | **UCSC genome browser plot for CNV breakpoints in TSPAN8 (hg19).** CNV breakpoints determined by cnvHitSeq for 1KG NGS data. Additional annotation for cnvHap derived breakpoints in NFBC 1986, Illumina Cardio-Metabochip probe locations and other publically available published CNVs breakpoints are also marked. Bottom section shows significant eQTL results from the GTeX project. Of note, significant eQTLs are tissue specific and are concentrated in the downstream regions near TSPAN8 exon11 and CNVR5583.1. **b** | **Frequency of TSPAN8 CNVs.** CNV deletion frequency (cnvHitSeq) of TSPAN8 exon 11 and CNVR5583.1 in different worldwide populations from the thousand genomes sequence data. **c** | **Linkage disequilibrium.** Heatmap of LD in the NFBC 1986 cohort (genotyped on the Illumina Cardio-Metabochip) for TSPAN8 exon 11 CNV deletion and the B-allele of SNP located within CNVR5583.1.

### Metabolomic signatures of TSPAN8 variants

Metabolomic signatures were obtained by applying univariate and multivariate approaches (Multiphen, see methods) using cnvHap derived CNV genotypes. Across TSPAN8 and within a window of one megabase around TSPAN8, the strongest CNV-metabolome association signal was discovered within TSPAN8 exon 11 (chr12:71523134), closely followed by exon 10 and other nearby probes (**Figure 2c, 2d, 2e**). At chr12:71523134, on meta-analysis (inverse variance fixed effects) of 228 metabolic phenotypes in NFBC 1986 and NFBC 1966 (n=9,190) we found the top metabolic phenotype to be **XL_HDL_TG** (Triglycerides in very large HDL, P=3.65 × 10^−4^). Genome-wide univariate inflation factors for CNV-**XL_HDL_TG** associations were found to be 0.97 and 1.18 in NFBC 1986 and NFBC 1966 respectively (**Supplementary Table ST 1a**). In our multivariate signature analysis, a different subclass of HDL (**XL_HDL_PL**: Phospholipids in very large HDL, P=0.000112) was found to be associated with multivariate joint signature P-value of 2.99 × 10^−11^ **Supplementary Table ST 1c vi**). Using intensity (LRR) based association model (association independent of cnvHap genotypes or CNV calling) **XL_HDL_TG** replicated in the meta-analysis of NFBC 1986 and NFBC 1966 with P value 0.039 (**Supplementary table ST 1d iii**). In a separate British replication cohort (whitehall) the strongest lipid association signal in the TSPAN8 gene was observed for HDL lipid at 12:71526064, near exon 10 with univariate LRR Pvalue=5.02 × 10^−6^, **Supplementary Table ST 1g i, Extended data figure 9a**).

**Figure 2.**
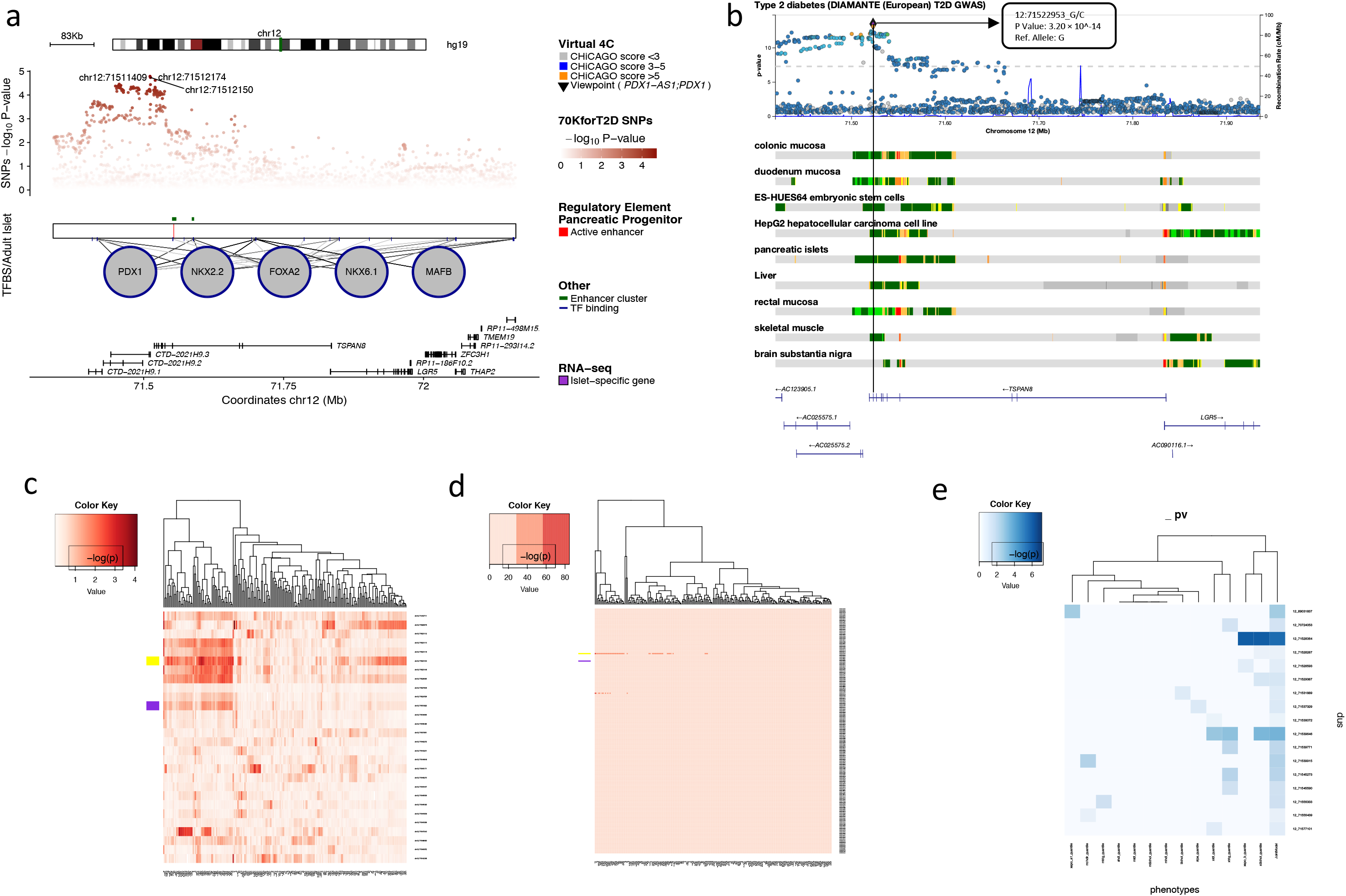
**a** | **Epigenetic annotation and T2D trait association.** Transcription factor binding sites (TFBS) in TSPAN8 determined in the adult human pancreatic islets. Schematic includes 70KforT2D SNP association results for type 2 diabetes along with regulatory annotations for TSPAN8 (negative HiC results). This schematic was generated by the Islet regulome browser, url: http://isletregulome.org/isletregulome/ with following settings: *Chromatin maps*: Pancreatic progenitors (**Cebola L, et al. 2015**), Enhancer clustering algorithm: *Enhancer clusters (Pasquali L, et al 2014)*, transcription factors: *Adult islets – Tissue specific (Pasquali L, et al 2014*) and SNP association results**: *70KforT2D.*** **b** | **Regional Manhattan plot.** SNP association results for TSPAN8 in T2D DIAMANTE GWAS analysis (N~ 1 million) obtained from the T2D knowledge portal (url: http://www.type2diabetesgenetics.org/). Schematic includes epigenetic annotations for transcriptional activity within TSPAN8. Of note, within a window of one megabase, the most significant SNP association locus for T2D-outcome lies near TSPAN8 exon 11 (rs1796330, chr12:71522953, P=3.20 × 10^−14^). **c** | **Metabolomic signatures in NFBC 1986.** Heatmap showing association results for TSPAN8 CNV genotypes with 227 metabolomic measurements in NFBC 1986. TSPAN8 exon 11 deletion showing high degree of pleiotropy, is marked in yellow and the T2D CNV deletion reported by the Wellcome Trust CNV study-*CNVR5583.1* is marked in violet. **d** | **Metabolomic signatures in NFBC 1986 and NFBC 1966.** Heatmap showing association results for TSPAN8 CNV genotypes with 227 metabolomic measurements in NFBC 1986 and NFBC 1966 pooled together (N~9000) within a window of one megabase around TSPAN8. **e** | **Multivariate metabolomic signature in whitehall cohort.** Heatmap showing association results for Log R ratio (LRR) with lipid phenotypes in Whitehall cohort. Association analysis was done using reverse regression. In this approach all phenotypes were analysed in a single joint model yielding a single P value for all phenotypes.

Further, CNV at chr12:71523134 (MAF~2%) exhibited strong pleiotropy with >50% (115/228) of the metabolites having a significant P Value<0.05 (**Figure 3a**). In contrast, significant SNP association results at the same position (MAF=43%) showed pleiotropy of only 2%. In addition individuals with CNV deletion at chr12:71523134 had significantly higher levels of metabolite levels, particularly for LDL and its subcategories (**Supplementary Table 1b**). We found 61% (22/36) of LDL and its subclasses had a significant P-value <0.05 for higher metabolite levels.

**Figure 3.**
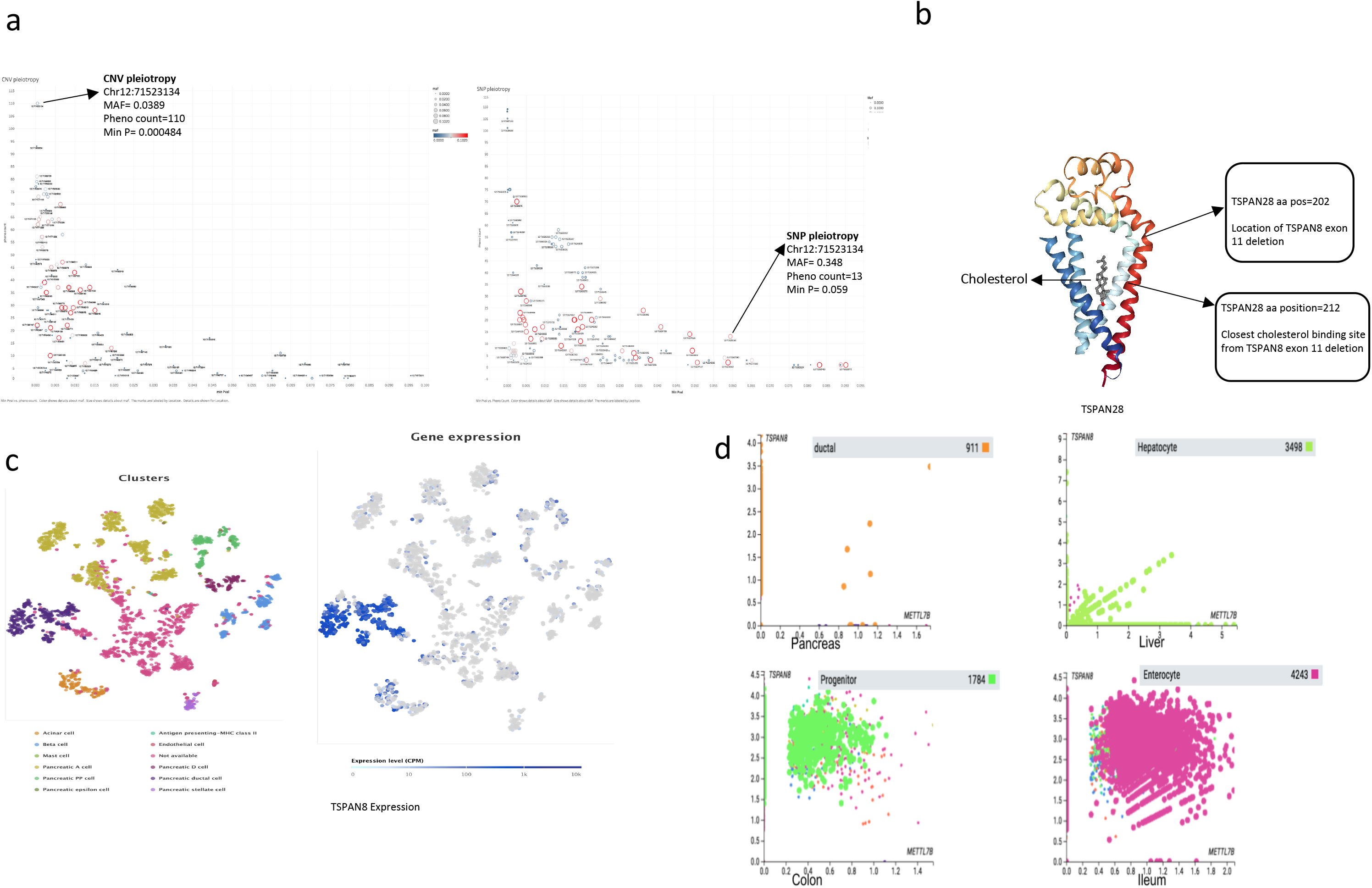
**a | Pleiotropic nature of TSPAN8 CNVs.** Schematic showing pleiotropic nature of TSPAN8 CNVs (left) and TSPAN8 SNPs (right) in NFBC 1986. Every probe location in the TSPAN8 gene is denoted by a circle and represents univariate association results for cnvHap genotypes with metabolomic measurements. Size and colour of each circle correspond to CNV allele frequency. Higher allele frequency is denoted by both colour (blue to red) and size of the circle. Y-axis denotes phenotype count i.e. number of metabolomic phenotypes found associated with given CNV genotype at a P value threshold of ~0.05. X-axis denotes the minimum P value observed at a given probe location. **b** | **3D protein structure of TSPAN28.** 3D protein structure from protein data bank showing a cholesterol-binding pocket in TSPAN28 (PDB id: **5TCX**). Through amino acid sequence alignment, TSPAN8 exon 11 deletion and its closest cholesterol-binding site have been mapped and highlighted. **c** | **Single-cell gene expression data in human pancreas.** Published single-cell gene expression data showing tissue-specificity of TSPAN8 in the ductal cells of human pancreas (**Single Cell Gene Expression Atlas**, Segerstolpe Å, Palasantza A et al. (2016) Single-Cell Transcriptome Profiling of Human Pancreatic Islets in Health and Type 2 Diabetes.). **d** | **Single-cell gene expression results from Human Cell Atlas.** Correlation of of single-cell gene expression data for TSPAN8 and METTL7B in specific cell types, namely: ductal cells in pancreas, hepatocytes in liver, progenitor cells in colon and Enterocyte in Illeum. Single-cell gene expression were obtained from the human cell atlas project and analysed and visualized through the cellxgene software (https://data.humancellatlas.org/analyze/portals/cellxgene)

We report all univariate and multivariate CNV genotype association and validation results for all common and rare CNVs in the TSPAN8 gene with 228 NMR characterized metabolomic phenotypes in NFBC 1986 and NFBC 1966 in **Supplementary Tables: ST 1b, ST 1c, ST 1d, ST 1e** and their subsections. We highlight two metabolites of interest from previous GWAS of human metabolome^15,16^ which lie near TSPAN8 exon 10 namely **1)** Ratio of 7-methylguanine to mannose (chr12:71524858, P=6.58 × 10^−7^) and **2)** 3,4-dihydroxybutyrate (chr12:71526064, P=7.36 × 10^−5^ synonym: 3,4-Dihydroxybutyric acid) (**Supplementary Table 1j**).

### Functional characterization and biology of TSPAN8

To understand the function of TSPAN8 CNVs further, we carried out genome wide TSPAN8 CNV-QTL (germline CNVs association with iPS cell line gene expression) analysis in human induced pluripotent stem cells from the HipSci project ^6^. One of the top hits included METTL7B (genome-wide rank 3, P=0.000195, Qvalue=0.865, **Supplementary Table 3b**). We observe that in addition to these results it might be possible to assign a-priori assumption for functional relationship between TSPAN8 and METTL7B based on current knowledge of common transcriptional factors, gene and protein co-expression and Phewas analysis. This a-priori link between the two genes is discussed in later sections and also reviewed separately in **Supplementary note**.

NKX2-2 is a common epigenetic regulator for TSPAN8 and METTL7B active in adult human pancreatic islets (**Supplementary Table 3a, Figure 2a, Extended data figure 7a**). Main evidence of tissue specific expression for these three genes included GTEx and Novartis whole body gene expression maps^17^ (**Extended data figures: ED 7d, ED 7e and ED 9f**), significant tissue specific SNP-eQTLs (GTEx, **Supplementary Table 3c**), eQTL colocalization and causality results reported by the T2D knowledge portal (**Supplementary Table 3e**), single-cell gene expression databases (**Supplementary Tables: ST 3f, ST 3g, ST 3h and ST 3i**) and whole body gene expression results in mouse (Tabula Muris, **Extended data Figure 4**), Papio anubis, Ovis aries and Xenopus laevis (**Supplementary Table ST 3o**). In developing Xenopus laevis, NKX2-2 and TSPAN8 expression seemed to have a positive correlation from Nieuwkoop and Faber stage^18^ (NF) stage 12 to 35-36 but from NF stage 35/36 they become negatively correlated, suggesting additional transcriptional repression factors in play. Such factors are unknown at the moment and warrants further investigation (**Extended data Figure 10**).

Thus combining evidence from multiple databases and publications we have demonstrated strong evidence of in-tandem RNA and protein co-expression for NKX2-2, TSPAN8 and METTL7B. We further hypothesize that these three genes together are likely to be functionally linked in energy homeostasis and glucose metabolism in the body through their coordinated action in tissues and organs related to insulin, hormones, other signaling molecule-**1)** production: pancreas **2)** processing: liver and gut **3)** regulation: brain and central nervous system and 4) uptake: muscles.

The protein structure of TSPAN8 remains unknown at the moment. However, the protein structure of TSPAN28 which has strong sequence similarity with TSPAN8 (E-value = 2 × 10^−38^) has been determined and further shown to bind with cholesterol^19^ (**Figure 3b**). Here, we show that TSPAN8 exon 11 deletion when mapped to TSPAN28 is about ten amino acids away from the closest cholesterol binding pocket, thus opening up the possibility that TSPAN8 exon 11 deletion might have an effect on cellular cholesterol transport, binding and metabolism.

### Disease, lifestyle and exposome analysis of TSPAN8, NKX2-2 and METTL7B

Here, we report novel CNV association with disease outcome for TSPAN8, NKX2-2 and METTL7B in previously characterized cohorts namely DESIR for T2D^20^, child obesity cohort^21^ and adult obesity cohort^21^ (**Supplementary Table 1f**).

LRR based association P values for these cohorts for the TSPAN8 exon 11 (chr12:7152314) were 0.0651, 0.00305 and 2.57 × 10^−12^ respectively. These results included several novel loci with significant CNV-disease signals.

In NFBC 1986 and NFBC 1966 the top phewas traits associated with TSPAN8, METTL7B and NKX2-2 included insulin medication, glycemic traits and smoking (**Supplementary Tables: ST 4a, ST 4b**); while common phenotypes from the FINNGEN project included- type 2 diabetes, diabetes with coma (both type 1 and 2), neurological complications and several categories of glycemic traits (**Supplementary Table 4d**). Some common lifestyle and exposome phenotypes from several public databases and GWAS catalogues included-death at home, medication use, BMI, hip circumference, waist hip ratio in females and balding pattern in males (**Extended data figure 3b**; **Supplementary Tables: ST 4c, ST 4e, ST 4f, ST 4g, ST 4h, ST 4i, ST 4j**). A common metabolomic signature for CNVs in TSPAN8, NKX2-2 and METTL7B included **XXL_VLDL_L (Total lipids in chylomicrons and extremely large VLDL)** which was recently reported to be associated with increased all-cause mortality rate in humans ^22^. In cancer biology, TSPAN8 has been well characterized and is mainly implicated in cancer hallmarks related to metastasis and angiogenesis. By comparing and contrasting mutations including CNVs, single nucleotide variants (SNVs), and gene expression, with a well known classic tumour suppressor gene like PTEN (**Extended data Figures: ED 8b, ED 8c**), we propose that TSPAN8 is likely to be a oncogene, involved with cancer metabolism through CNV amplifications and overexpression. Thus, since TSPAN8 SNVs are quite sparse, TSPAN8 CNVs are more likely to be cancer driver events. Overall survival estimates for patients with overexpression in TSPAN8 in many cancer types was also found to be significantly lower (**Supplementary Table 4k**).

## Discussion

Finnish populations are known to be enriched for deleterious variants, hence are likely to be of added value for understanding molecular mechanisms of common disease such as type 2 diabetes and metabolic disorders. Here, we have reported one of the first in-depth association analyses of CNVs using univariate and multivariate approaches in TSPAN8 gene with 228 circulating plasma metabolites in over 9,300 Finnish individuals. In our analysis we have highlighted some important aspects related to CNV detection and association approaches for cohorts with large sample sizes, commonly characterized through microarrays and NGS platforms. Some salient points included successful application of ‘population aware’ methods for CNV detection, application of probabilistic measures for CNV genotypes for improved CNV-phenotype associations and leveraging intensity based approaches for independent validation of CNV-phenotype associations. We demonstrate that CNVs are prevalent in germline, somatic and iPS cell line genomes, their characterization, especially determining correct breakpoints and allele frequency remain challenging and underexplored. Importantly, approaches for delineating functional impact of bystander CNVs from real disease causing pathogenic variants remain limited at the moment. New technologies based on CRISPR based genome engineering, long read sequencing and sequence guided reanalysis of published GWAS microarray datasets, are some promising leads to address some of these challenges.

Using a modest sample size of ~9,100 our multivariate approach of using all 228 metabolomic phenotypes in a single model allowed us to pinpoint the most significant and also perhaps the functionally important region in TSPAN8 located within exon 11. In contrast, the multi-ethnic DIAMANTE meta-analysis for type 2 diabetes ^23^ reported the most significant SNP in TSPAN8 near exon 10 at 12:71522953 with P Value=10^−14^ using a sample size of ~ 1 million (74,124 T2D cases and 824,006 controls). This result highlights the power of multivariate metabolomics analysis for genomics and highlights its relevance for rare variant analysis which usually require extremely large sample sizes.

In the HipSci data, we rediscovered TSPAN8 CNV deletions in iPS donor genomes with MAF=5% and subsequent CNV-QTL analysis led to the discovery of METTL7B, as a potential new trans CNV-QTL for TSPAN8. Further NKX2-2 was found to be a common transcription factor for these two genes active in pancreatic islets. The initial evidence from the iPS cell lines analysis is suggestive but weak, due to non-significant Qvalue, however several additional results from epigenetic and single-cell RNA-Seq data reinforced our hypothesis that TSPAN8, METTL7B and NKX2-2 are likely to be functionally linked in a very tissue-specific manner in humans and other species. This evidence enabled us to build a robust a-priori hypothesis for TSPAN8 and METTL7B and give more weight to the HipSci results. An additional important result we would like to highlight is the possible involvement of TSPAN8, METTL7B and NKX2-2 in the PPAR pathway. To elucidate 1) NKX2-2 has been experimentally shown to regulate TSPAN8 and PPARG (**Supplementary Table 3a**) and 2) the fact that METTL7B (Synonym: ALDI-Associated With Lipid Droplets 1^24^) physically co-localizes with PLIN1 on peroxisomes in the cell cytosol, suggests that the PPAR pathway might indeed be a common denominator (**Extended data figure 7b**).

Another observation we make here is that in addition to strong tissue-specific gene-expression in humans and other species, TSPAN8, NKX2-2 and METTL7B further tend to be expressed in pairs but never together i.e. all three genes being expressed in the same tissue is rarely seen. This phenomenon of pair-exclusivity of gene expression was also indirectly reflected through Kaplan-Meier survival curve estimates for many cancer types (**Supplementary Table 4k**). Some highlights of such patterns included pancreatic ductal carcinoma and kidney cancer, both of which have strong germline tissue expression. One exception to this pattern was cervical cancer where all three genes were found to be overexpressed. Cervical cancer has links to human papillomavirus (HPV), thus might be a genuine outlier. However, we caution that these observations are preliminary and require further experimental investigation before any definitive conclusions can be made.

TSPAN8 and METTL7B, both seem to have strong evidence of being involved with obesity. TSPAN8’s role in obesity is strongly indicated by knockout experiment in mice leading resistance to weight gain and also corroborated by our novel association results for child obesity, where we found deletions in TSPAN8 are protective against obesity with an odds ratio of 24.59 (Supplementary Table ST 2k i). METTL7B’s role in obesity is a relatively new observation. Of importance is a recent GWAS analysis of childhood onset obesity^25^ where the authors reported rs540249707 near METTL7B to have an odds ratio of 3.6 (95% CI 2.13-6.08, P=1.77 × 10^−6^) which was higher than the FTO variant rs9928094 with odds ratio=1.44 (95% CI 1.33-1.57, P=1.42 × 10^−18^). Further, METTL7B variants have also been reported as one of the top hits in GWAS of amphetamine response (**Supplementary Table 4f ii**). Though discontinued, amphetamines are known to be prescribed as anti-obesity medication^26^ with side effects related to increased alertness. Whether association of TSPAN8 and METTL7B with obesity, central nervous system or other traits is driven by independent molecular mechanisms or through common molecular pathways is left unvalidated at the moment.

Findings from single-cell data indicate that TSPAN8 is mainly expressed in pancreatic ductal and acinar cells, thus highlighting its involvement of the neuro-exocrine axis for energy homeostasis and metabolism. Further, NKX2-2 and TSPAN8 seem to be strongly co-expressed in similar regions of the human brain, in particular in the midbrain region around hypothalamus, neural stem and spinal cord. METTL7B on the other hand is overexpressed in glioblastoma. Observations of several fold high expressions for TSPAN8 and NKX2-2 are replicated in the UK Brain expression study^27^ (**Extended data figure 6**) and were also reflected through results from MetaXcan, eCAVIAR and COLOC analysis for TSPAN8 (**Supplementary Table 3e**). These observations for TSPAN8 and NKX2-2 constitute one of the first reported genetic links in the neuro-exocrine axis for energy and metabolic homeostasis in humans. Our neurological observations are further intriguing due to an earlier reported association of TSPAN8 SNPs in exon 10 with 3,4 dihydroxybutyrate (synonym: 3,4-Dihydroxybutyric acid). Butyrate has hormone-like properties and can induce enhanced secretion of glucagon and insulin ^28^ in the pancreas and has known beneficial effects on intestinal homeostasis for energy metabolism via the gut-brain axis^29^. Importantly, 3,4-Dihydroxybutyric acid is known to be linked to satiety^30,31^ and with ultra-rare Succinic Semialdehyde Dehydrogenase Deficiency (SSADH).

Using principles similar to reverse genetics, through Phewas and phenotypic trait analysis, we further strengthen our metabolomic and gene expression findings. One such example is a common metabolomic signature for TSPAN8, METTL7B and NKX2-2 CNVs-**XXL_VLDL_L**, which was recently found to be associated with all-cause mortality ^22^. The mortality risk factor is further corroborated by strong Phewas signal for traits related to death at home in UK Biobank results which were common for all three genes. Some of the other phenotypic traits of interest included high medication use, diabetes with neurological complications, several categories of glycemic traits, BMI, hip-circumference and fat mass.

Our observation from current scientific literature is that all common germline CNV deletions (Atleast 5 CNVs with MAF>5%) in TSPAN8 are nearly depleted in almost all somatic cancer genomes. The fact that they are also depleted in iPS cell line genomes postulates that TSPAN8 CNVs are likely to be under unknown somatic evolutionary forces. In contrast, genes like GSTM1 or RHD which also harbour common germline CNV deletions with MAF>30% seem to retain CNV deletions during their somatic evolution (data not presented). This phenomenon indicates that human germline genomes might have inbuilt safety mechanisms or harbour tumour suppressive variants, in order to provide inherent protection against uncontrolled cell proliferation or cancer. Like TSPAN8 CNV deletions, one might expect such tumour suppressive events to be present as ‘common variants’ in various human populations.

Of note, germline metabolomic signatures of TSPAN8 and its associated genes can shed light on cancer metabolism, which can be exploited for diagnostic, therapeutic or palliative interventions. One such possibility which warrants further investigation is our observation that TSPAN8 and METTL7B (active but with weak expression) is expressed with high specificity in triple-negative breast cancer (**Extended data figure 5**).

To conclude, our results robustly demonstrate the strong pleiotropic effects of TSPAN8, METTL7B and NKX2-2 on a wide range of human phenotypes, suggesting common molecular mechanisms and biological pathways, which opens up new possibilities for diagnostic and therapeutic approaches for metabolic diseases.

## Methods

### Study cohorts

All cohorts reported in this study including data from 1KG project, NFBC 1986, NFBC 1966, Whitehall II study (WH-II), DESIR, Child Obesity cohort, Adult Obesity cohort, 1958 British Birth Cohort 1958 (BC1958), National Blood Survey (NBS), Helsinki Birth cohort (HBCS) and HipSci samples have prior ethical approval and consent from all study subjects involved. Further details including aims and methods have been reported earlier. In our analysis we refer to NFBC 1986 as the primary discovery cohort for CNVs and metabolomic signatures and NFBC 1966 and WH-II as replication cohorts. BC1958, NBS and HBCS were used as control cohorts for ascertaining CNV allele frequencies. Child obesity, Adult obesity and DESIR cohorts were used for replicating disease outcomes. 1KG NGS data was used for CNV breakpoint and frequency calculations. HipSci data was used for functional characterization of CNVs.

### Metabonomic measurements in NFBC 1986 and NFBC 1966

Metabolomic measurements for NFBC 1986 (n=228) and NFBC 1966 (n=228) cohorts were carried out using high throughput ^1^H Nuclear Magnetic Resonance (NMR) technology developed by Nightingale Healthcare Limited.

Further details of aims and methods for characterization of various lipoprotein species, ratios, size along with other metabolites have been described earlier. A complete list of metabonomic phenotypes used in our analysis, their names and categories are listed in **Supplementary Table 5**.

### Lipid measurements in WHII

After obtaining relevant permissions, we had access to the following lipid measurements for our analysis-Apoprotein A1 (Apo A1), Apoprotein B (Apo B), Cholesterol total (Bchol), Cholesterol HDL (HDL), Intermediate density lipoprotein (IDL), Triglycerides (Trig), Lipoprotein A (LPA) and Cholesterol LDL (LDL). Additional details related to the WH-II study and their phenotypes are available from the consortium website (https://www.ucl.ac.uk/whitehallII/).

### CNV Analysis

#### NGS based CNV identification

First, CNV calls were generated using the cnvHiTSeq algorithm^32^ in TSPAN8 genic region using NGS low coverage data from 1KG project for 17 different populations. cnvHiTSeq uses a Hidden Markov Model (HMM) based probabilistic model for genotyping and discovering CNVs from NGS platforms. It incorporates various signatures from sequencing data such as read depth, read pair and allele frequency information and then integrates them into a single HMM model to provide improved sensitivity for CNV detection. Normalisation of the sequence data prior to CNV analysis using cnvHitSeq was performed in the following manner-sequencing files in binary alignment format (bam) for the different populations were first downloaded from the 1KG website. For each population, samples were normalised in a sliding window of 25 base pairs and were corrected for wave effects and GC content. Next, cnvHiTSeq was run with a combination of read depth and split read information, with an initial transition probability of 0.15 and 15 expectation maximization (EM) training iterations.

### cnvHap - Normalisation and quality control

Cohorts genotyped on the Illumina platform were processed through the Illumina Beadstudio (now called Genomestudio2.0) software. Log R Ratio (LRR), B-Allele frequency (BAF) and sample SNP genotypes were exported from the Beadstudio software as ‘final reports’ for subsequent CNV analysis. Prior to CNV calling, data normalisation was done in a genotyping plate specific manner in order to correct for batch effects. For every genotyping plate, data was adjusted for LRR median correction and LRR variance. Genomic wave effects were accounted for by fitting a localised loess function with a 500 kbp window. Next, the processed LRR and BAF values with relevant covariates such as genotyping plate, BAF, LRR variance etc were used as input by the cnvHap software for CNV calling.

### CNV predictions using cnvHap

CNVs in the TSPAN8 gene were called in various cohorts using the cnvHap algorithm^33^. This algorithm uses a haplotype hidden Markov model (HMM) model for simultaneously discovering and genotyping CNVs from various high-throughput SNP genotyping platforms such as Illumina CardioMetabochip and Agilent aCGH arrays. The haplotype HMM of cnvHap uses combined information of CNVs (LRR) and SNP (BAF) data in population aware mode for CNV predictions. cnvHap has specific emission parameters for different genotyping platforms. In our analysis, we used Illumina platform specific emission parameters in all cases. cnvHap was used in its population aware mode where all samples were simultaneously used to train the model. In contrast, CNV detection methods such as PennCNV trains its HMM model one sample at a time and does not leverage population level information for CNV prediction.

### CNV segmentation

cnvHap calculates the most probable linear sequence of copy number states (Hidden state of the HMM model) for each sample by using dynamic programming and outputs this sequence as CNV breakpoints. Additionally, cnvHap also calculates probabilistic CNV genotypes or expected CNV genotypes (described later) based on posterior probabilities. Of note, CNV allele frequency based on breakpoints information might differ from frequency calculated using posterior probabilities of CNV genotypes.

### Expected CNV genotypes

The haplotype HMM of cnvHap calculates the probability of deletion and duplication for each sample at a given probe which we refer to as the expected CNV genotypes. For example, at a particular probe if a sample has CNV genotype assigned as 1 (heterozygous deletion) with probability of 0.8 then the expected CNV genotype is calculated as

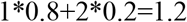

The expected CNV genotypes were calculated separately for deletions and duplications for every sample and at every probe location. We have used expected CNV genotypes for all our association analysis and results.

### Association analysis

Next, using the MultiPhen software^34^ we carried out both univariate and multivariate approaches for associating expected CNV genotypes with metabonomic phenotypes in all cohorts. In the univariate analysis, for every probe location P Values for association was calculated using expected CNV genotypes as predictors and metabonomic phenotypes as the outcome. For common genomic probe locations, meta-analysis of NFBC 1986 and NFBC 1966 was performed using the inverse variance fixed-effect model.

For multivariate analysis, we used the MultiPhen software which implements a reverse regression model where phenotypes are used as predictors and CNV genotypes are used as outcome. We refer to this model as a multivariate joint model. The multivariate joint model uses ordinal probit regression to associate CNV genotypes (outcome) with multiple metabolomic phenotypes (predictors) simultaneously and provides a single joint P Value capturing the effect of all phenotypes together. In addition, we have further implemented a variable selection method into this model by using a custom backward-selection algorithm. This backward-selection method reduces the correlation structure in the phenotypic space through an iterative process and in the end provides a set of uncorrelated variables. This uncorrelated set of variables is next used in the standard MultiPhen multivariate joint model to obtain a single P value and effect size for all phenotypes. In our analysis we refer to this subset of phenotypes as metabolomic signature. In all univariate and multivariate regression analyses, phenotypes were transformed using quantile normalisation and 50 LRR principal components (PCs), LRR variance and sex were used as covariates

### Intensity based validation of CNV association

There have been several reports regarding the use of direct raw signal data from various technology platforms without using intermediate processing or software as an input for bioinformatics methods. Such approaches have previously been applied for CNV-phenotype association studies where LRR intensity measurements from genotyping platforms were used^35^. Here, we have leveraged a similar approach by using LRR-phenotype association results as an alternate method to validate CNV genotype-phenotype association results. Similar to CNV association analysis, we applied univariate and multivariate approaches from MultiPhen for the LRR data and used for 50 LRR PCs, LRR variance and sex as covariates in the model.

### Multiple Testing

Previous studies have reported the presence of high degree of correlation in metabolomic and lipid phenotypes. In order to adjust for multiple testing thresholds in the presence of such correlation structure, several alternate methods to Bonferroni correction such as the Sidak-Nyholt correction have been proposed ^36^. Briefly in this method, for calculating the net number of effective tests M_eff_ in the presence of correlation structure in the phenotypes, the following formula can be used.

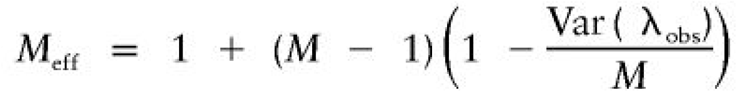

Here lambda_obs_ is the eigen decomposition of the correlation matrix of metabonomic phenotypes. The net effective number of tests M_eff_ obtained can then be applied to the Sidak formula or the Bonferroni correction, in order to determine the correct P Value threshold. On applying this correction to Sidak-Nyholt and the Bonferroni method, the adjusted multiple testing the P value thresholds obtained were 8.05 × 10^−4^ and 7.8×10^−4^ respectively.

### Linkage disequilibrium

In NFBC 1986 and other cohorts LD calculation was done using pearson correlation coefficient for LRR and CNV genotype data from genotyping arrays and sequence data. In addition, a linear regression model was also used to calculate LD between CNV-genotype and the number of B-alleles. In NFBC 1966 no probes were found to be in LD (r^2^>0.5) with TSPAN8 exon 11 or CNVR5583.1, hence not reported.

## Supporting information

Supplemental Table Index

## ACKNOWLEDGEMENTS

We would like to thank Luise Cederkvist Kristiansen for preliminary work and analysis of NFBC cohorts.

## URLS

http://mips.helmholtz-muenchen.de/proj/GWAS/gwas/

http://isletregulome.org/isletregulome/

https://www.proteinatlas.org/

https://gtexportal.org/home/

https://rdrr.io/cran/MultiPhen/

http://r2.finngen.fi/

https://atlas.ctglab.nl/

http://www.type2diabetesgenetics.org/home/portalHome

https://www.targetvalidation.org/

https://cancer.sanger.ac.uk/cosmic

https://www.cbioportal.org/

https://www.ebi.ac.uk/gwas/

http://www.phenoscanner.medschl.cam.ac.uk/

https://www.imperial.ac.uk/people/l.coin

https://portal.brain-map.org/

http://www.braineac.org/

https://www.internationalgenome.org/

https://www.oulu.fi/nfbc/

https://www.ucl.ac.uk/epidemiology-health-care/research/epidemiology-and-public-health/research/whitehall-ii

http://www.xenbase.org/entry/

https://www.gsea-msigdb.org/gsea/msigdb/annotate.jsp

https://www.ebi.ac.uk/gxa/sc/home

http://www.hipsci.org/

https://genome.ucsc.edu/

https://www.wtccc.org.uk/wtcccplus_cnv/supplemental.html

https://bioinfo.uth.edu/scrnaseqdb/

https://pax-db.org/

https://www.ebi.ac.uk/arrayexpress/experiments/E-MTAB-2836/

http://www.nealelab.is/uk-biobank

http://phewas.mrbase.org/

http://geneatlas.roslin.ed.ac.uk/

https://kmplot.com/analysis/

## Life Sciences Reporting Summary

Further information on experimental design is available in the Life Sciences Reporting Summary.

## Code availability

Code used to process and analyze data is available at Multiphen: https://github.com/lachlancoin/MultiPhen cnvHap: https://www.imperial.ac.uk/people/l.coin cnvHitSeq: https://sourceforge.net/projects/cnvhitseq/

## Data and material availability

NFBC data and material: https://www.oulu.fi/nfbc/materialrequest

## AUTHOR CONTRIBUTIONS

**T.D, LJMC, MRJ, MRJ** Involved in study design, performed analysis and wrote the manuscript.

**AG, D.G.** Carried out analysis of HipSci data and helped in writing the manuscript. **D.S.** Advised on statistical inference and helped in writing the manuscript, **P.F.** Provided access and advised on obesity and type 2 diabetes cohorts.

## COMPETING FINANCIAL INTERESTS

None

## ADDITIONAL INFORMATION

1. Extended data figures: ED 1 - ED 10.
2. Supplementary Tables: ST 1 - ST 5
3. Supplementary Information contains: Legends for Supplementary Tables ST1-ST5, Supplementary note and Single-cell data from Human Cell Atlas.

## Materials & Correspondence

Should be addressed to T.D.

---

**Table 1.** Biology of TSPAN8. Overview of TSPAN8 biology, insights and corresponding evidence.

| TSPAN8 biology | Status | Evidence |
| --- | --- | --- |
| Common CNVs in TSPAN8 and their frequency in worldwide populations | New results | Supplementary tables: ST 2c, ST 2d, ST 2e, ST 2g |
| Association and replication of TSPAN8 variants with various metabolites and its effect on LDL | New results and insights | Supplementary tables: ST 1b, ST 1c, ST 1e, ST 1g, ST 1h,ST 1j |
| Pleiotropic nature of TSPAN8 variants (CNV & SNP) and allele frequency | New results | Fig 3a |
| Validation of CNV-phenotype association through an independent model | New results | Supplementary table: ST 1d |
| Association of TSPAN8 loci with mortality, diabetes, diabetes with coma,obesity and 3,4-dihydroxybutyrate (satiety, SSADH) | New results and insights | Supplementary tables: ST 1f, ST 1j, ST 4d, ST 4h |
| TSPAN8 CNVs and CNV-eQTL in Human induced pluripotent stem cells | New results | Extended data: ED Fig. 7c; Supplementary table: ST 3b |
| Direction of causality and lifestyle effects of NKX2-2, TSPAN8 and METTL7B variants | New results | Extended data: ED Fig. 3b |
| Cloest cholesterol binding site to TSPAN8 exon 11 deletion | New insights | Fig. 3b |
| Knockout of TSPAN8 in mice | New insights | Champy, M., Le Voci, L., Selloum, M. et al. Reduced body weight in male Tspan8-deficient mice. Int J Obes 35, 605–617 (2011). |
| Role of TSPAN8 in PPAR pathway | New results | Extended data: ED Fig 3b, ED Fig 7b; Supplementary table: ST 3a |
| GTeX eQTLs for NKX2-2, TSPAN8 and METTL7B in different tissues of the human body | New insights | Supplementary table: ST 3c |
| Gene expression profile of TSPAN8 in various regions of the human brain and central nervous system | New insights | Extended data: ED Fig 6; Supplementary table: ST 3e |
| Transcriptional and epigenetic regulation of TSPAN8, METTL7B and NKX2-2 in different tissues | New insights | Fig 2b, Fig 7a; Supplementary tables: ST 3n |
| Role of TSPAN8 and NKX2-2 in pancreatic and organ development in model organisms Xenopus laevis | New insights | Supplementary table: ST 3o |
| Single cell gene expression of NKX2-2, TSPAN8 and METTL7B in 20 mouse organs in Tabula Muris | New insights | Extended data: ED Fig 4 |
| Single-cell expression profile of TSPAN8 in triple-negative breast caner | New insights | Extended data: ED Fig 5 |
| RNA, protein co-expression of NKX2-2, TSPAN8 and METTL7B in human tissues | New insights | Extended data: ED 7d, ED 7e; Supplementary tables: ST 3f, ST3g, ST 3h, ST 3i, ST 3j, ST 3k, ST 3l, ST3m, ST 3o |
| CNV profile of TSPAN8 in cancer genomes | New insights | Extended data: ED Fig 8a, ED Fig 8c |
| Gene expression profile of TSPAN8 in cancer transcriptomes and effect on cancer survival (Kaplan-Meier estimates) | New insights | Extended data: ED Fig 8b, ED 8c, Supplementary table: ST 4k |
| Gender effects on TSPAN8 function and association results | New results | Supplementary tables: ST 1g, ST 3l, ST 4c (iv, v, vi), ST 3o (Ovis aries, male and female seperately) |
| Total protein abundance of NKX2-2, TSPAN8 and METTL7B in the tree of life | New insights | Supplementary table: ST 3m |
| Therapeutic potential of TSPAN8 | New results | Supplementary table: ST 4b (keyword=insulin medication), ST 4h (keyword=treatment) |

**Extended data Figure 1.**
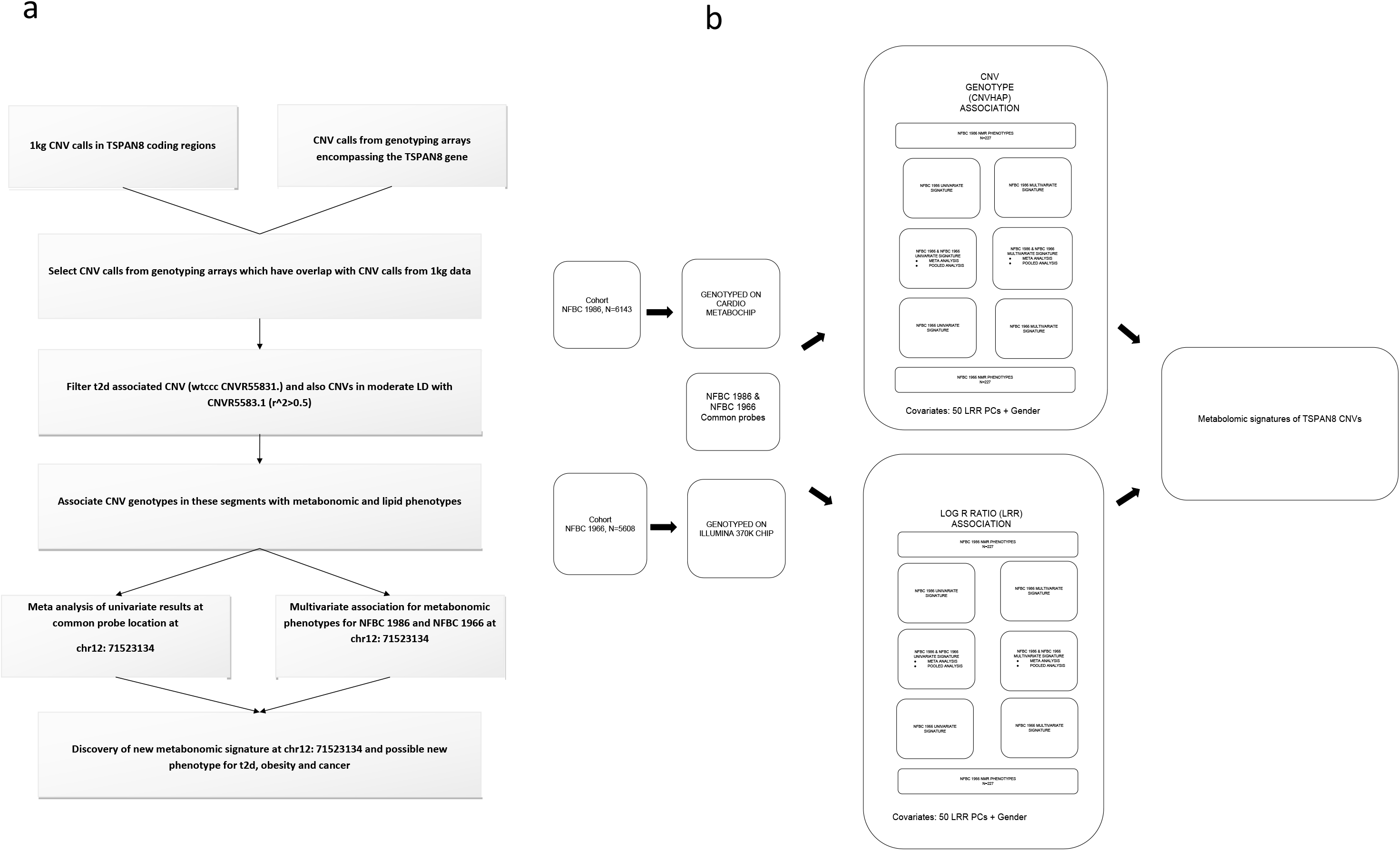
**a** | **TSPAN8 exon 11 signatures** Flow diagram showing steps leading to the determination of metabolomic signatures and a new phenotype for TSPAN8 exon 11 deletion. **b** | **Overview of association approaches** Association approaches using MultiPhen software for associating a) CNV genotypes and b) LRR with 227 metabolomic phenotypes in NFBC 1986 and NFBC 1966. Methods include both univariate, meta-analysis and multivariate approaches.

**Extended data Figure 2.**
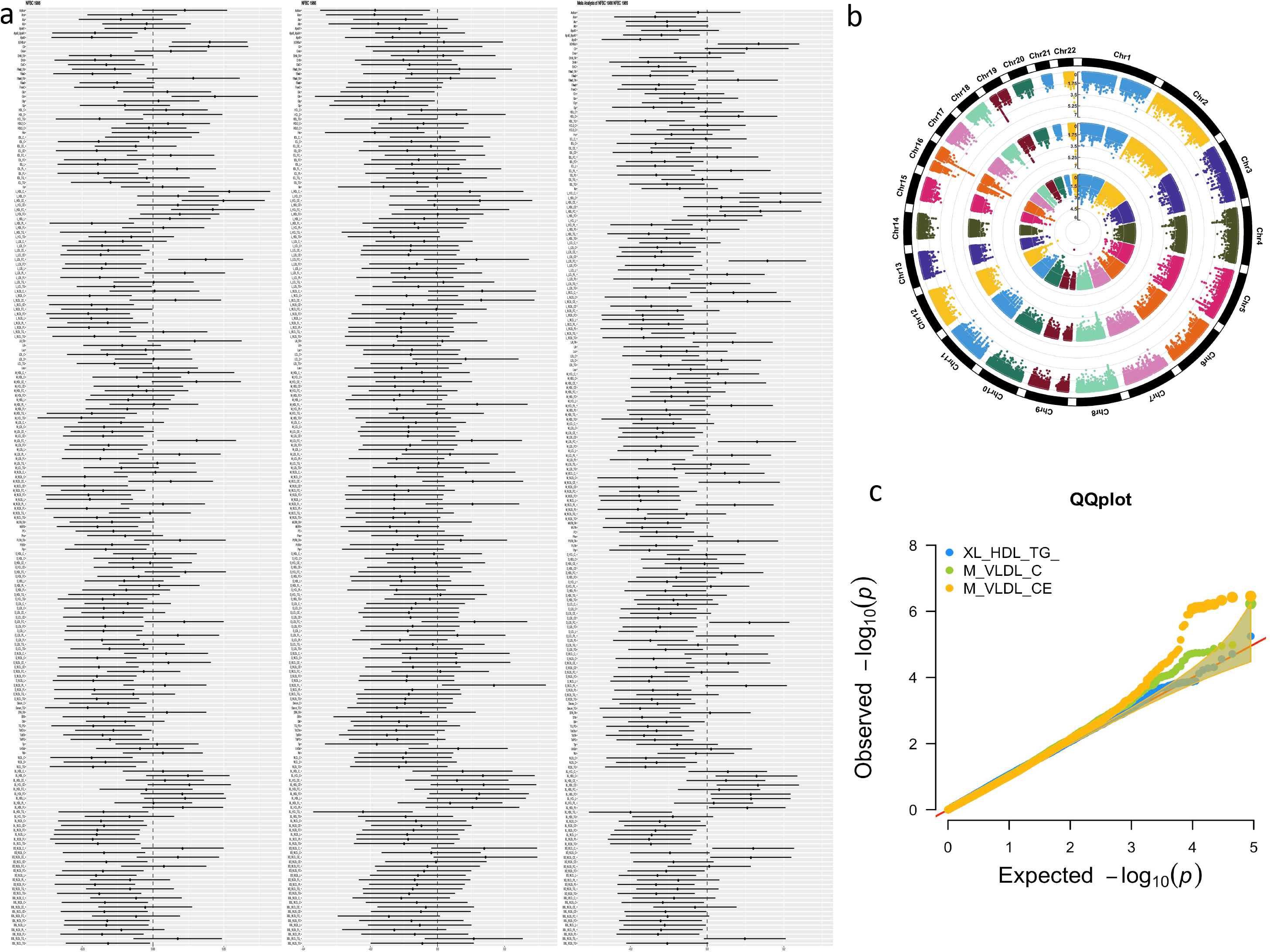
**a** | **Forrest plot** Forrest plot of beta coefficients and confidence intervals for CNV-phenotype association results at chr12:71523134 (TSPAN8 exon 11) in NFBC 1986, NFBC 1966 and meta-analysis of NFBC 1986 and NFBC 1966. **b** | **Genome-wide Manhattan plot** Circular Manhattan plot of genome-wide association results for the top three metabolomic phenotypes obtained on meta-analysis of 227 metabolomic phenotypes at chr12:71523134 (TSPAN8 exon 11). Order of phenotypes in the schematic-inner most circle to outer most circle: 1) **XL_HDL_TG_** 2) **M_VLDL_C** 3) **M_VLDL_CE.** **c** | **Quantile-Quantile plot** QQ plot for top three meta-analysis phenotypes at chr12:71523134 (TSPAN8 exon 11): 1) **XL_HDL_TG_** 2) **M_VLDL_C** 3) **M_VLDL_CE**

**Extended data figure 3.**
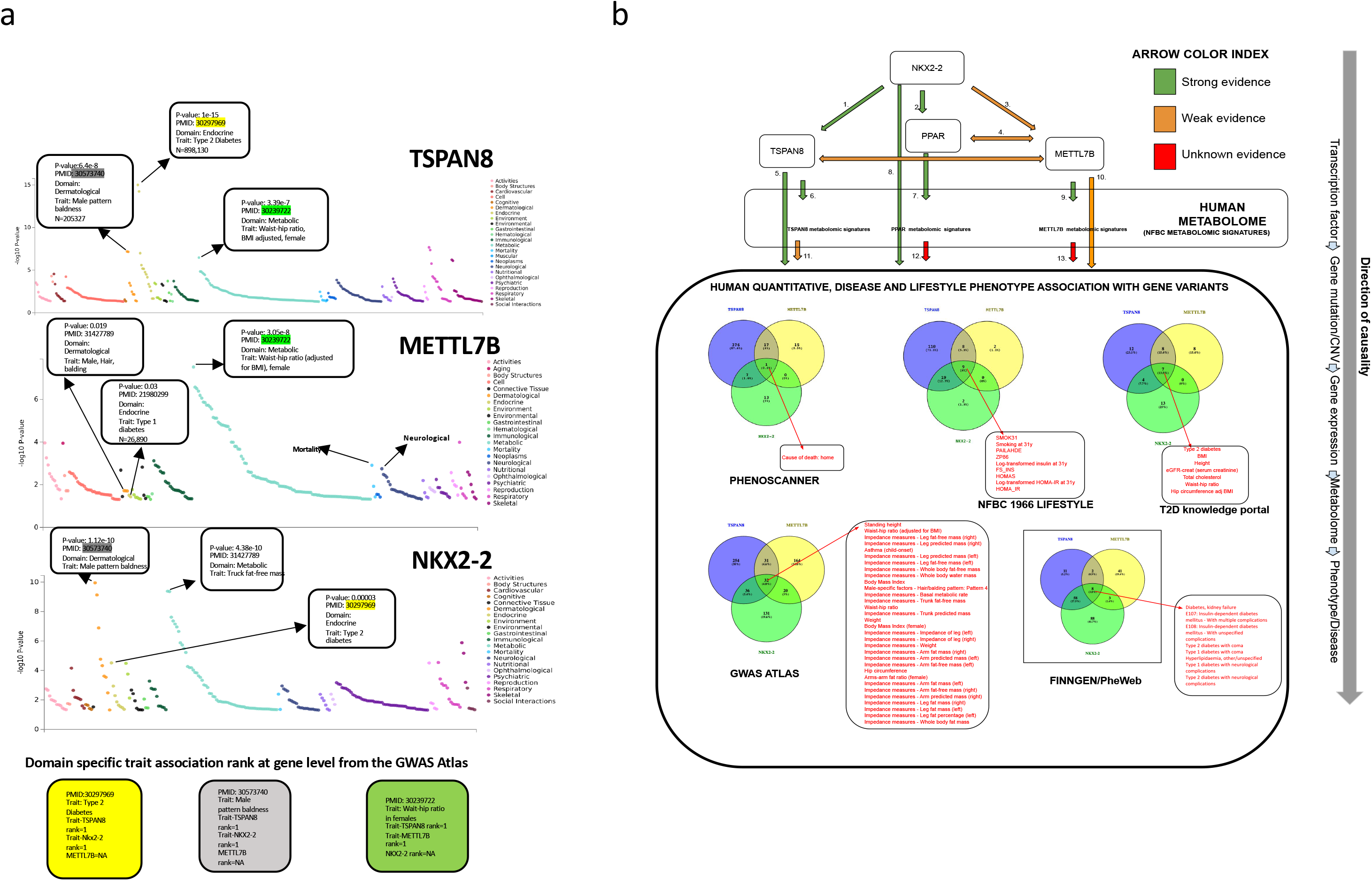
**a** | **GWAS-ATLAS Phewas** PheWas plot generated by the GWASATLAS server (https://atlas.ctglab.nl/PheWAS). Phenotypic traits associated with TSPAN8, NKX2-2 and METTL7B are sorted by their broader phenotypic domain and then P values. TSPAN8 shares the top trait with both NKX2-2 and METTL7B in metabolic, endocrine and dermatological domains and the top associated SNP reported is from same pubmed study. These results are based on 4756 GWAS studies (March 2020). **b** | **Direction of causality** Flow diagram outlining the direction of causality of NKX2-2 (common transcriptional factor), TSPAN8, METTL7B and PPARG pathway. In addition, various pieces of evidence showing how these genes influence with each other is annotated, including respective metabolomic signatures and common GWAS/PheWAS traits.

**Extended data figure 4.**
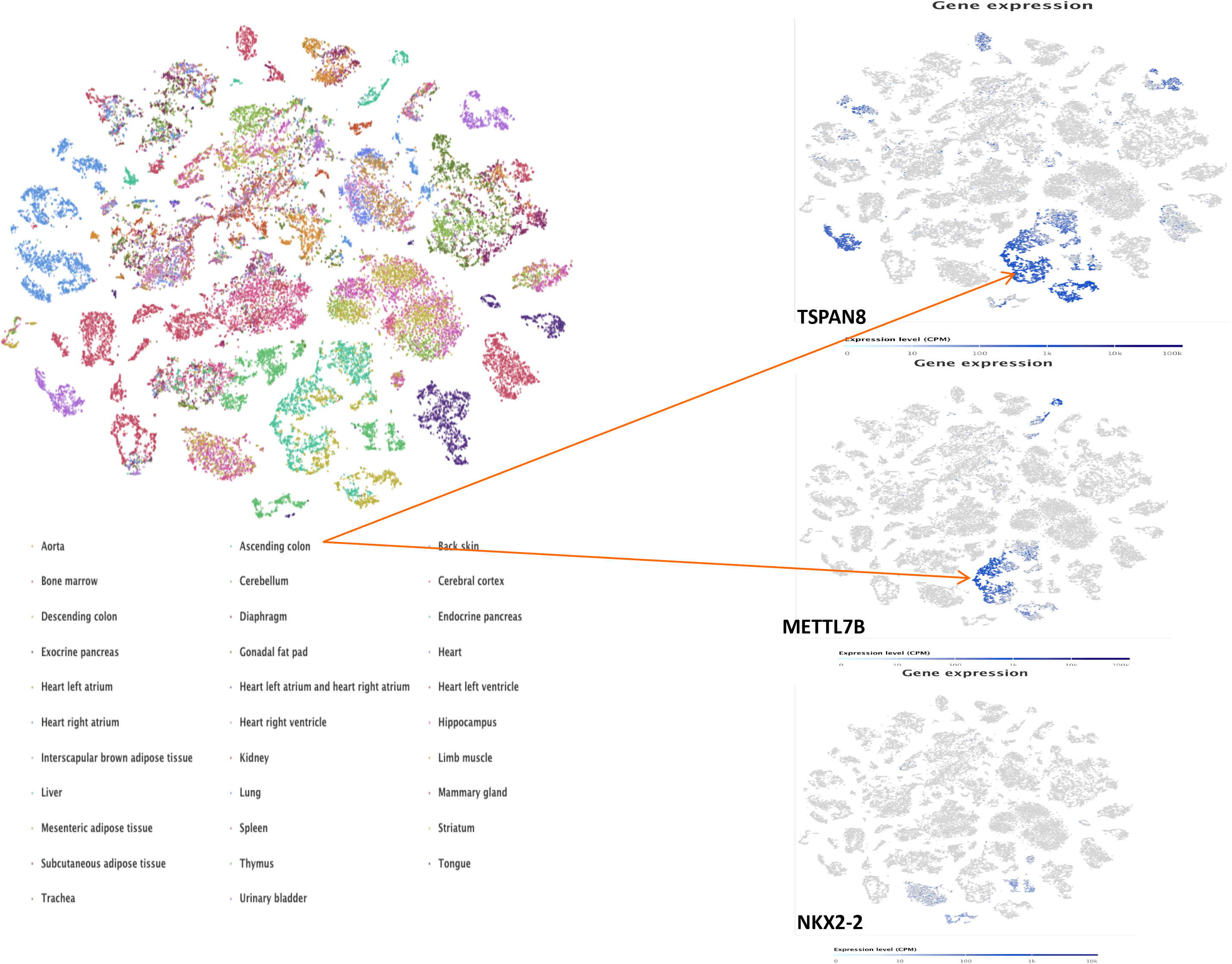
Single-cell RNA sequencing results from *Tabula muris*. Figure showing gene expression of TSPAN8, NKX2-2 and METTL7B in 20 different different organs in Tabula muris mice (Single Cell Gene Expression database, https://www.nature.com/articles/s41586-018-0590-4).

**Extended data figure 5.**
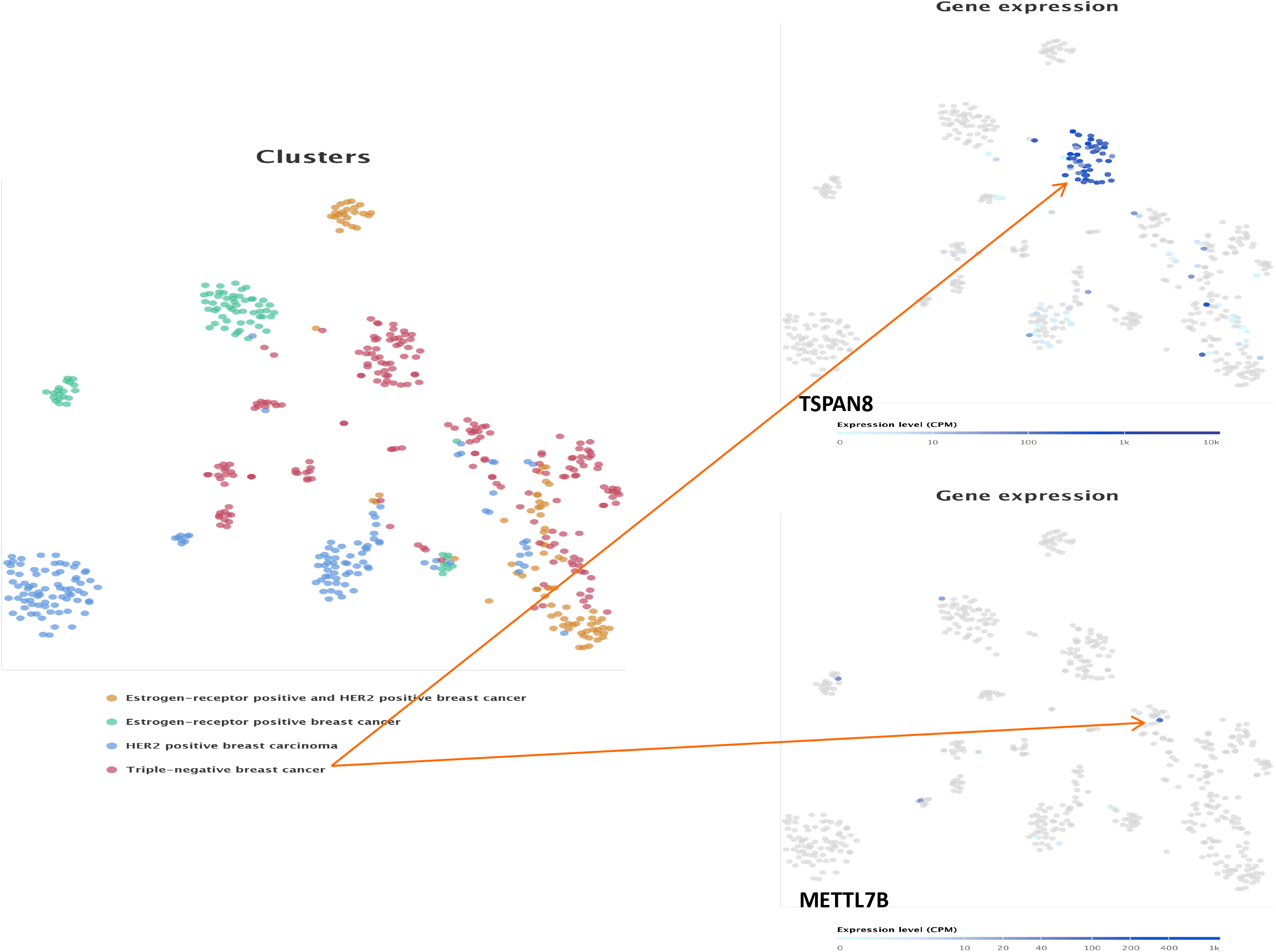
a | Single-cell RNA sequencing analysis from breast cancer subtypes. Gene expression levels of TSPAN8, NKX2-2 and METTL7B to different breast cancer types. TSPAN8 is strongly expressed in triple-negative breast cancer subtype. Strongest expression of METTL7B was also found to be in triple-negative breast cancer subtype but found only in one cell. (Single Cell Gene Expression database, Chung W et al. (2017) Single-cell RNA-seq enables comprehensive tumour and immune cell profiling in primary breast cancer)

**Extended data figure 6.**
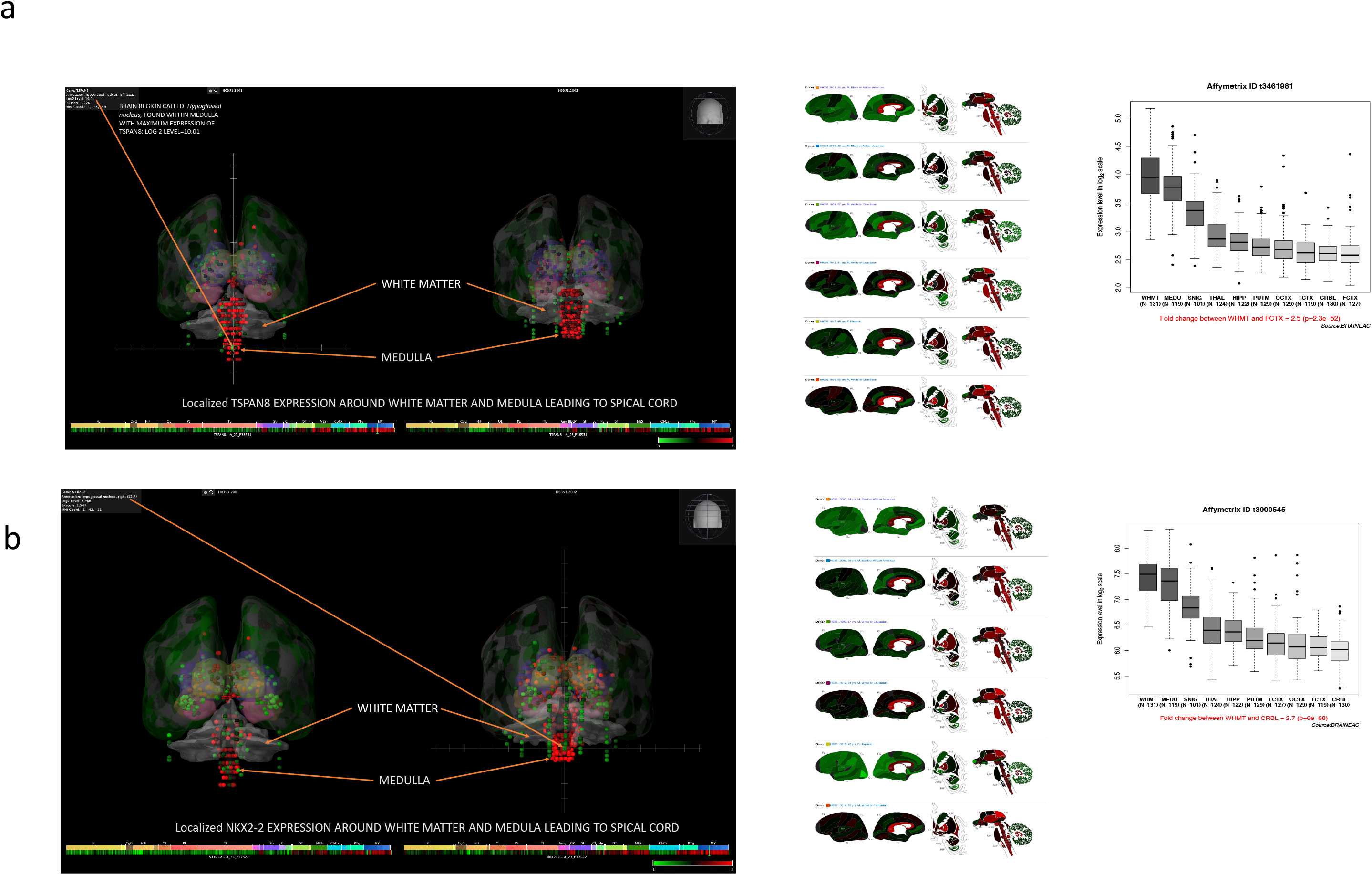
**a and 6b Gene expression results from the Allen Brain Atlas** **a**: **Left panel:** Annotation of gene expression of TSPAN8 on the 3D atlas of the human brain from the Allen brain consortium (https://portal.brain-map.org/). Left section and right section brains are from individuals *H0351.2001* and *H0351.2002* respectively. Maximum expression of TSPAN8 was found to be in the midbrain region, specifically white matter and medulla. In one individual high level of TSPAN8 expression in the Hypoglossal nucleus was *Log-2 level =10.01*. **Middle panel:** 2D lateral representation of the human brain from six individuals from Allen Brain Atlas portal showing the log2 level gene expression for TSPAN8. **Right panel:** TSPAN8 expression in ten regions of the human brain obtained from the BRAINEAC project (http://www.braineac.org/). Maximum expression of TSPAN8 white matter and medulla observed in the the Allen Brain Atlas is replicated in the BRAINEAC study. **b** | **Same as 6a but for NKX2-2**

**Extended data figure 7.**
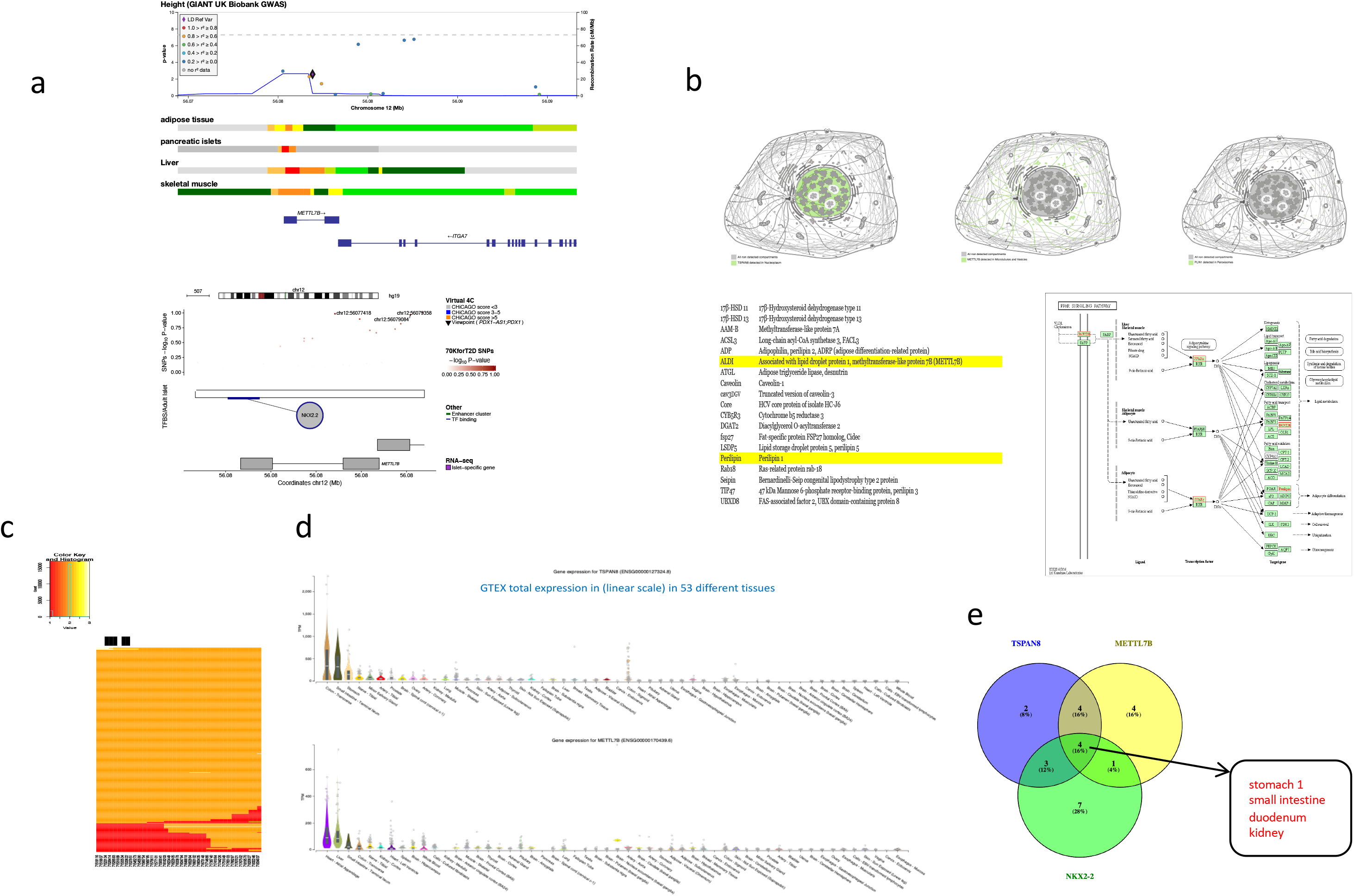
**Biological and functional insights for TSPAN8** **a:** Figure showing transcriptional factor binding activity (strong, weak etc) for METTL7B in various tissues of the human body obtained from T2D knowledge portal (http://www.type2diabetesgenetics.org/). Bottom figure confirms that NKX2-2 has a transcriptional factor binding role for METLL7B in adult human islets (http://isletregulome.org/isletregulome/). **b:** Upper panel: Subcellular location of TSPAN8, METTL7B and PLIN1 obtained from the human protein atlas (https://www.proteinatlas.org/). TSPAN8 is detected in the nucleoplasm where as METTL7B and PLIN1 are co located in the cell cytosol in microtubules, vesicles and peroxisomes. Bottom left: Published table confirming that ALDI (METTL7B) and PLIN1 are both lipid droplet proteins, confirmed by microscopy. **c** TSPAN8 CNVs detected in the germline HipSci donor samples. **d** Tissue specific gene expression for TSPAN8 and METTL7B obtained from the GTeX portal. **e:** Comparison of protein expression levels in various organs and tissues of the human body for TSPAN8, METTL7B and NKX2-2. Data obtained from the human protein atlas: https://www.proteinatlas.org/.

**Extended data figure 8.**
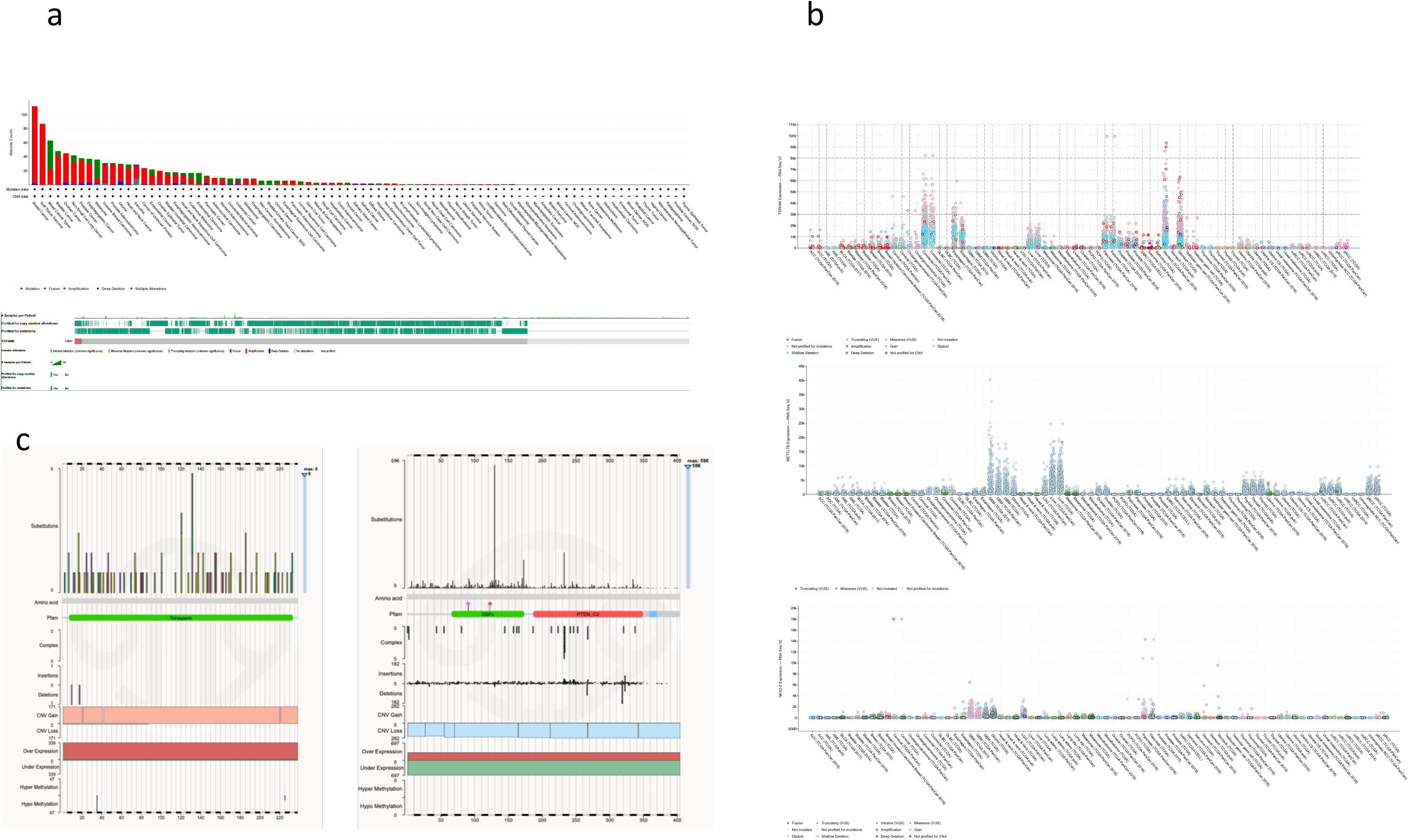
**Cancer genome and transcriptome analysis** **a** Landscape of mutations (CNV, Point mutations and indels) in ~70, 000 samples from the TCGA project in various cancer types. (https://www.cancer.gov/about-nci/organization/ccg/research/structural-genomics/tcga) TSPAN8 CNV amplifications is observed in about 1.8% of the samples along with negligible CNV deletions and point mutations. **b** Transcriptomic landscape of TSPAN8 in various cancer types from the TCGA project. **c** Comparison of mutations (CNV, SNVs and indels) and gene expression in TSPAN8 and PTEN from the COSMIC database. PTEN being a classic tumour suppressor has CNV deletions and an under-expression profile. TSPAN8 has an opposite mutation and gene expression profile with CNV amplifications and overexpression and no CNV deletions. This indicates TSPAN8 might have a oncogenic role in tumour biology.

**Extended data figure 9.**
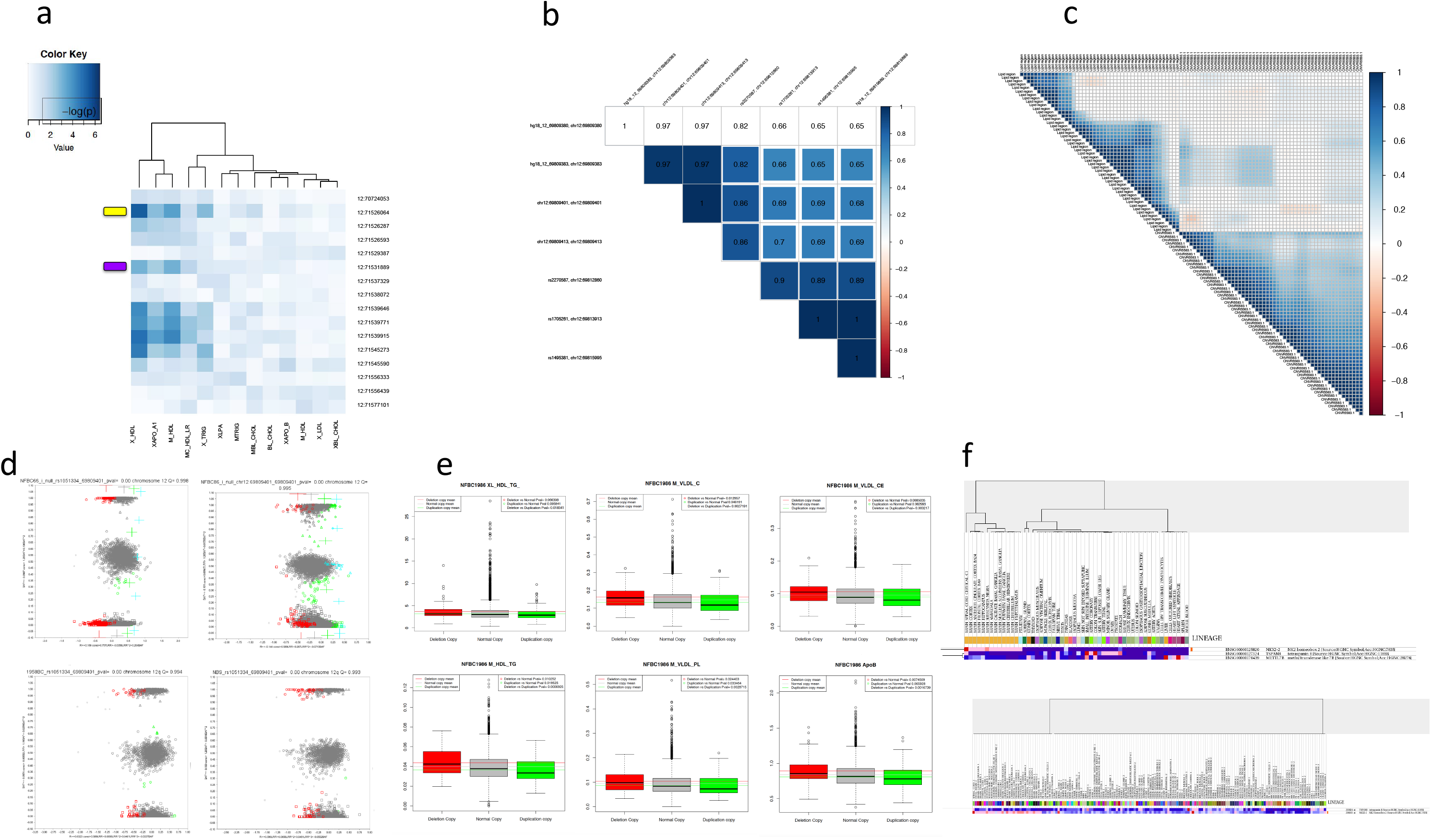
**Additional results** **a)** Univariate association results in the TSPAN8 gene region for Log R Ratio with various lipid species in the Whitehall cohort. Gender was not used as a covariate. Region nearest to the TSPAN8 exon 11 deletion is highlighted in yellow and region overlapping with CNVR5583.1 is highlighted in violet. Results with gender as a covariate is reported in Supplementary table **b)** Figure showing correlation (LD) between CNV genotypes in NFBC 1986 for exon 10 (rs2270587, chr12:69812860, build 36/hg18), exon 11 (chr12:69809401, chr12:69809401, build 36/hg18) and CNVR5583.1 (hg18_12_69819889, chr12:69819889, build 36/hg18). Original probe names are marked and genomic locations are based on human reference **genome build 36/hg18**. **c)** Figure showing the correlation (LD) of raw read depth values between TSPAN8 exon 11 CNV deletion (pleiotropic region) and CNVR5583.1, for the Finnish population from the thousand genomes project (FIN). **d)** Two-dimensional cluster plots plot for TSPAN8 exon 11 CNV in NFBC 1966 and NFBC 1986. × and Y axes in the plot corresponds to LRR and BAF respectively. Each point on the plot represents a sample; shape of the point refers to the number of B-alleles in the SNP genotype; colour represents CN state (pink=0, red=1, grey=2, green=3, light blue=4). Original probe names are marked and genomic locations are based on human reference **genome build 36/hg18**. **e)** Box plots for top six meta-analysis metabolomic phenotypes in NFBC 1986 individuals stratified by CNVs (deletion and duplication) at chr12:71523134 (TSPAN8 exon 11). Colour denoted are: Red=Individuals with CNV deletion, Grey=Individuals with Normal homozygous copy, Green=Individuals with CNV Duplication. Red, grey and green lines indicate the respective means of the metabonomic phenotypes for samples with the deletion, normal and duplication CNVs. Non-parametric association P values between the different groups are shown on the top right corner. **f)** Unsupervised clustering of gene expression of TSPAN8, NKX2-2 and METTL7B in different tissues of the human body from the a) GTeX project (top) and b) Novartis whole body gene expression map (bottom). The heatmap of expression values are denoted by the following colour scheme in the order of **highest to lowest gene expression levels**: *red, peach, pink, light blue, dark blue purple.* Of note, TSPAN8, NKX2-2 and METTL7B are expressed in pairs but never together in the same tissue. Interesting observations include **GTex**: 1) paired expression of NKX2-2 and TSPAN8 in spinal cord and 2) paired expression of TSPAN8 and METTL7B in small intestine and colon; **Novartis whole body gene expression map**: 1) paired expression of TSPAN8 and METTL7B in spinal cord 1 and 2 and pancreatic islet 1 and 2.

**Extended data figure 10.**
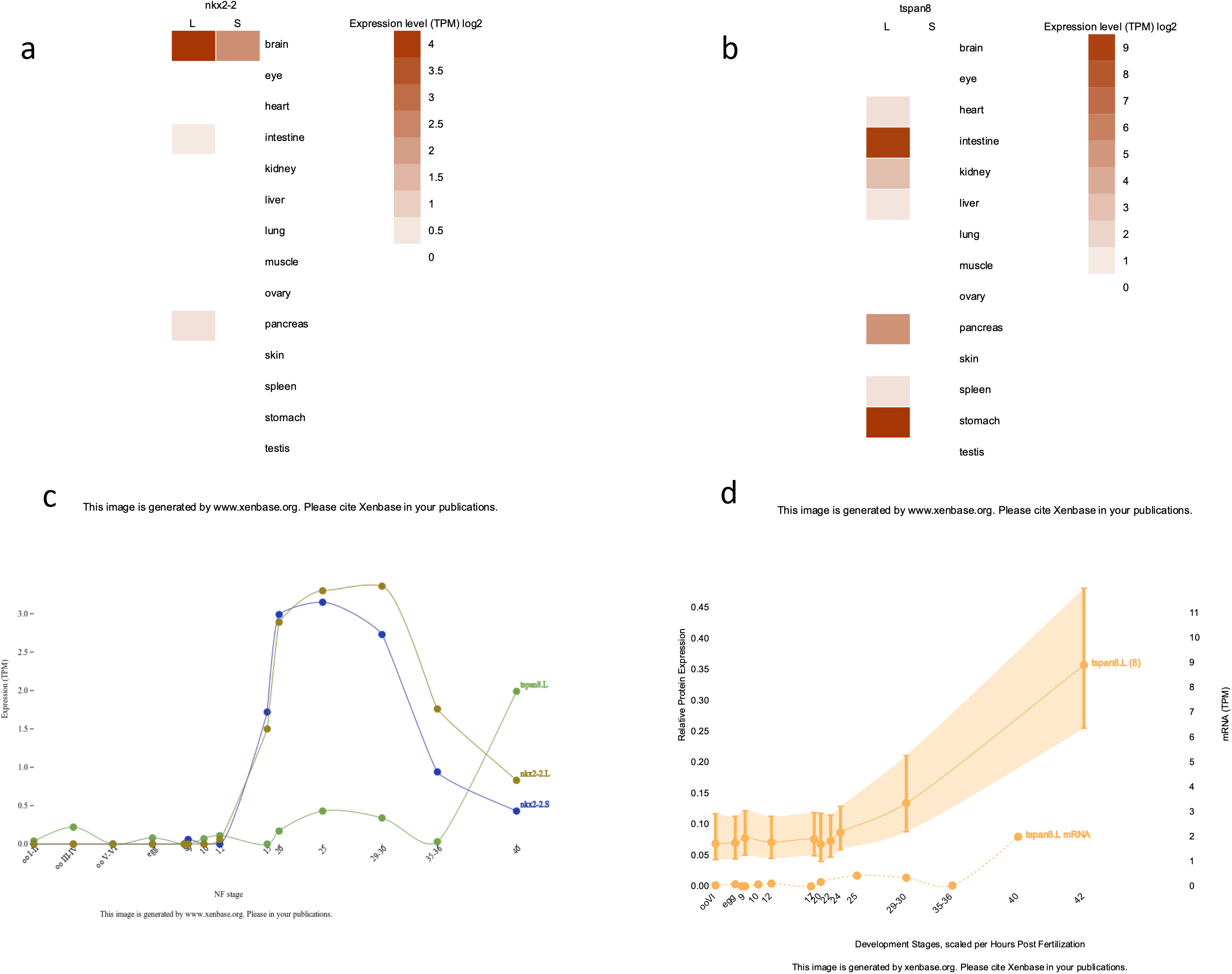
Results from Xenbase. **a)** RNA-Seq derived expression level (TPM) of nkx2-2 in different tissue of Xenopus laevis **b)** RNA-Seq derived expression level (TPM) of tspan8 in different tissue of Xenopus laevis **c)** Gene expression dynamics of different isoforms of tspan8 RNA and nkx2-2 during different developmental stages of Xenopus laevis **d)** Correlation of protein and mRNA levels of tspan8 during different developmental stages of Xenopus laevis

## Notes

### Competing Interest Statement

The authors have declared no competing interest.

